# Dsuite - fast D-statistics and related admixture evidence from VCF files

**DOI:** 10.1101/634477

**Authors:** Milan Malinsky, Michael Matschiner, Hannes Svardal

**Affiliations:** Zoological Institute, University of Basel, Basel, Switzerland; Department of Paleontology and Museum, University of Zurich, Zurich, Switzerland; Department of Biosciences, University of Oslo, Oslo, Norway; Department of Biology, University of Antwerp, Antwerp, Belgium; Naturalis Biodiversity Center, Leiden, The Netherlands

**Keywords:** ABBA-BABA, *D* statistic, f4-ratio, gene flow, introgression, software

## Abstract

1. Patterson’s *D*, also known as the ABBA-BABA statistic, and related statistics such as the *f*_4_-ratio, are commonly used to assess evidence of gene flow between populations or closely related species. Currently available implementations require custom file formats and are impractical to evaluate all gene flow hypotheses across datasets with many populations or species.
2. Dsuite is a fast C++ implementation, allowing genome scale calculations of the *D* and *f*_4_-ratio statistics across all combinations of tens or hundreds of populations or species directly from a variant call format (VCF) file. Furthermore, the program can provide evidence of whether introgression is confined to specific loci and aid in interpretation of a system of *f*_4_-ratio results by implementing the ‘f-branch’ method.
3. Dsuite is available at https://github.com/millanek/Dsuite, is straightforward to use, substantially more computationally efficient than other comparable programs, and presents a novel suite of tools and statistics, including some not previously available in any software package.
4. Thus, Dsuite facilitates assessment of evidence for gene flow, especially across large genomic datasets.

## Introduction

Admixture between populations and hybridization between species are common and a bifurcating tree is often insufficient to capture their evolutionary history (Green *et al.* 2010; Patterson *et al.* 2012; Tung & Barreiro 2017; Kozak *et al.* 2018; Malinsky *et al.* 2018). Patterson’s *D* statistic, first used to detect introgression between modern human and Neanderthal populations (Green *et al.* 2010; Durand *et al.* 2011), has since then been widely applied and used across a broad range of taxa (Fontaine *et al.* 2015; vonHoldt *et al.* 2016; Tung & Barreiro 2017; Kozak *et al.* 2018; Malinsky *et al.* 2018). The *D* statistic and the related estimate of admixture fraction *f*, referred to as the *f*_*4*_-ratio (Patterson *et al.* 2012), are simple to calculate and well suited for taking advantage of genomic-scale datasets, while being robust under most demographic scenarios (Durand *et al.* 2011).

Programs for calculating *D* and the *f*_*4*_-ratio from genomic data include ADMIXTOOLS (Patterson *et al.* 2012), HyDe (Blischak *et al.* 2018), and Comp-D (Mussmann *et al.* 2019). However, what limits their utility is that none of these programs can handle the variant call format (VCF) (Danecek *et al.* 2011), the standard file format for storing genetic polymorphism data produced by variant callers such as samtools (Li 2011) and GATK (DePristo *et al.* 2011). Moreover, as each calculation of *D* and *f* applies to four populations or taxa, the number of calculations/quartets grows rapidly with the size of the dataset. The number of quartets is 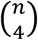, i.e. *n* choose 4, where *n* is the number of populations. This presents challenges both in terms of increased computational requirements and for interpretation of the results. It is partly for these reasons that previous studies utilizing *D* and the *f*_4_-ratio involved small numbers of populations or taxa, with few exceptions (Kozak *et al.* 2018; Malinsky *et al.* 2018). With more genomic data becoming available, there is a need for handling datasets with tens and up to hundreds of taxa. Dsuite addresses the above issues in that it calculates *D* and *f*_4_-ratio statistics directly from VCF files, is substantially more efficient than other programs, and provides an implementation of the *f*-branch statistic (Malinsky *et al.* 2018) to aid interpretation. Finally, unlike the other software packages, Dsuite calculates statistics specifically designed to investigate signatures of introgression in genomic windows along chromosomes.

## Methods and implementation

The *D* and *f*_4_-ratio statistics calculated by Dsuite are usually presented as applying to biallelic SNPs across four populations or taxa: P1, P2, P3, and O, related by the rooted tree (((P1,P2),P3),O), where the outgroup O defines the ancestral allele, denoted by A, and the derived allele is denoted by B (Green *et al.* 2010; Durand *et al.* 2011; Pease & Hahn 2015). The site patterns are ordered such that the pattern BBAA refers to P1 and P2 sharing the derived allele, ABBA to P2 and P3 sharing the derived allele, and BABA to P1 and P3 sharing the derived allele. Under the null hypothesis, which assumes no gene flow, the ABBA and BABA patterns are expected to occur with equal frequencies, and a significant deviation from that expectation is consistent with introgression between P3 and either P1 or P2. See especially (Durand *et al.* 2011) for more detail.

While simple site pattern counts can be computed for single sequences, the Dsuite implementation works with allele frequency estimates, so multiple individuals can, and ideally should, be included from each population or taxon. Denoting the derived allele frequency estimate at site *i* in P1 as 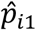 the following sums are calculated across all *n* biallelic sites:

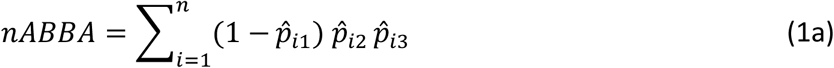

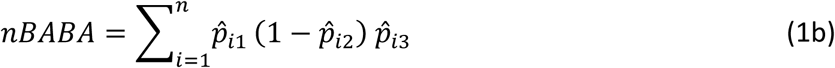

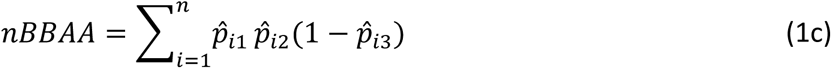

in cases where 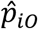, the derived allele frequency in the outgroup, is equal to zero.

### The Dtrios program

Dsuite does not assume *a priori* knowledge of population or species relationships, only the outgroup has to be specified. The first subprogram, `Dtrios` calculates the sums in equation (1) for all trios of populations or taxa in the dataset. The command produces three types of output. For the first, in a file with the “BBAA.txt” suffix, `Dtrios` attempts to infer the population or species relationships: it orders each trio assuming that the correct tree is the one where the BBAA pattern is more common than the discordant ABBA and BABA patterns, which are assumed to result for example from incomplete lineage sorting, repeated mutation at the same site, or from introgression. In addition, P1 and P2 are ordered so that *nABBA* >= *nBABA* and, therefore

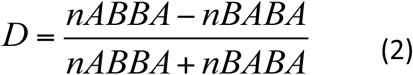

is never negative. The second type of output is the *D*_min_ score, the minimum *D* for each trio regardless of any assumptions about the tree topology (Malinsky *et al.* 2018). There is no attempt to infer the true tree; instead, the trio is ordered so that the difference between *nABBA* and *nBABA* is minimized. This output is in a file with the “Dmin.txt” suffix. Finally, there is also an option for the user to supply a tree in Newick format specifying known or hypothesized relationships between the populations or species. An output file with the “tree.txt” suffix then contains *D* and *f*_4_-ratio values for trios ordered in a way consistent with this tree.

Where the frequency of the derived allele in the outgroup is not zero, the results of `Dtrios` correspond to the *D* and *f*_4_-ratio statistics as defined by Patterson *et al.* (2012), who present the statistics as applying to an unrooted four taxon tree, with O being simply a fourth population rather than an outgroup. Their D definition is:

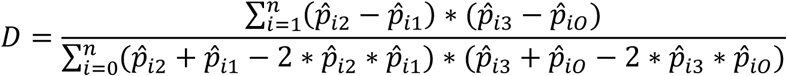

In this case, the ancestral vs. derived allele assignment is not necessary and the A and B labels can be assigned arbitrarily; the BAAB site pattern is equivalent to ABBA, ABAB to BABA, and AABB to BBAA. Therefore, the Patterson *et al.* (2012) definition of *D* corresponds to changing the right-hand side of equations (1a - c) to:

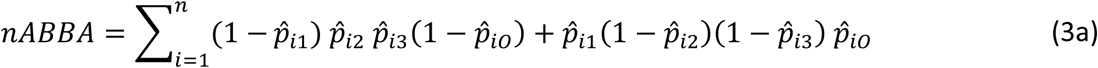

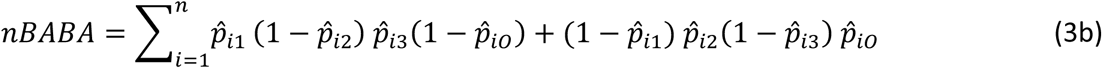

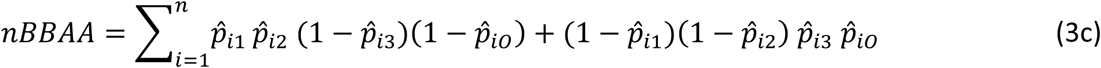

Thus, the *D* statistic can in principle be calculated without an outgroup. However, in designing Dsuite, we assume that an outgroup is usually available, which reduces the complexity of the analysis and of downstream interpretation of the results.

To assess whether *D* is significantly different from zero, `Dtrios` uses a standard block-jackknife procedure as in Green *et al.* (2010) and Durand *et al.* (2011), obtaining an approximately normally distributed standard error. For all three types of output, `Dtrios` calculates the Z-scores as *Z* = *D*/*std*_*err*(*D*), and outputs the associated p-values. However, when testing more than one trio, users should take into account the multiple testing problem and adjust the p-values accordingly. Although the different *D* statistics calculated on the same dataset are not independent, a straightforward conservative approach is considering them as such and controlling for overall false discovery rate.

Calculating the *f*_4_-ratio requires that P3 be split into two subsets, P3a and P3b, which is done in Dsuite by randomly sampling alleles from P3 at each SNP and is possible even if the dataset contains only one individual from P3. The results then follow the Patterson *et al*. (2012) definition:

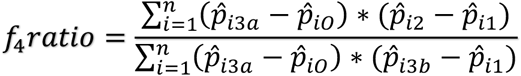

### The DtriosCombine program

It is common practice, especially for larger datasets, that VCF files are divided into smaller subsets by genomic regions, e.g. per chromosome. This facilitates the parallelization of many computational workflows. The `DtriosCombine` program enables parallel computation of the *D* and *f*_4_-ratio statistics across genomic regions, by combining the outputs of multiple `Dtrios` runs, summing up the counts in equations (3a - c) and the denominator of the *f*_4_-ratio. It also calculates overall block-jackknife standard error across all regions to produce overall combined p-values for the *D* statistic.

### The Dinvestigate program

The program `Dinvestigate` provides further information about trios for which the *D* statistic is significantly different from zero by calculating *f*_d_ (Martin *et al.* 2015) and *f*_dM_ (Malinsky *et al.* 2015) in sliding genomic windows. These statistics are specifically suited for application to genomic windows and can be used to assess whether the admixture signal is confined to specific loci and to assist in locating any such loci. For each trio specified by the user, the program outputs overall *f*_d_, and *f*_dM_, and also produces a text file which contains the values of *f*_d_ and *f*_dM_ in sliding windows. The size of the windows is specified by the user and refers to a fixed number of ‘informative’ SNPs, i.e. SNPs that change the numerator of these statistics for any particular trio.

### The Fbranch program

The number of possible gene flow donor-recipient combinations increases rapidly in datasets with more than four populations or taxa. A unified test for introgression has been developed for a five taxon symmetric phylogeny, implemented in the D_FOIL_ package (Pease & Hahn 2015), however, no such framework currently exists for datasets with six or more taxa. A common approach is to perform the *D* and *f*_4_-ratio analyses on four taxon subsamples from the dataset (e.g. (Green *et al.* 2010; Martin *et al.* 2013; vonHoldt *et al.* 2016; Kozak *et al.* 2018; Malinsky *et al.* 2018). However, the number of analyses that need to be performed grows very quickly. Given a fixed outgroup, the number of combinations is 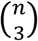, i.e. *n* choose 3, where *n* is the number of taxa. For example, there are 1,140 different combinations of ((P1, P2), P3) in a dataset of 20 taxa, growing to 161,700 combinations in a dataset with 100 taxa. Interpreting the results of such a system of four taxon tests is not straightforward; the different subsets are not independent as soon as the taxa share drift (that is, they share branches on the phylogeny) and, therefore, a single gene flow event can be responsible for many elevated *D* and *f*_4_-ratio results. At the same time, the correlations, especially of the *f*_4_-ratio scores, can be informative about the timing and the donor-recipient combinations for introgression events.

The f-branch or *f*_*b*_(C) metric was introduced in Malinsky *et al.* (2018) to disentangle correlated *f*_4_-ratio results and assign gene flow evidence to specific, possibly internal, branches on a phylogeny by building upon the logic developed by Martin *et al.* (2013). It is implemented in Dsuite `Fbranch`, to our knowledge the first publicly available implementation of this statistic. Given a specific tree (with known or hypothesized relationships), the *f*_*b*_(C) statistic reflects excess sharing of alleles between the population or taxon C and the descendants of the branch labelled b, relative to allele sharing between C and the descendants of the sister branch of b. Formally:

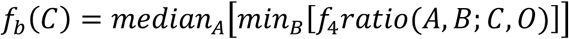

where B refers to the populations or taxa descending from the branch *b*, and *A* refers to descendants from the sister branch of *b*. The calculation is over all *f*_4_-ratio results which had A and B in either the P1 or P2 positions and C in the P3 position. The phylogenetic tree for `Fbranch` should be supplied in Newick format and should be the same tree as used in the `Dtrios` and/or `DtriosCombine` calculations.

## Performance and results

We assessed the performance of Dsuite using three datasets: 1) variants mapping to the largest *Metriaclima zebra* reference genome scaffold (∼16Mb) from a dataset comprising 73 species of Lake Malawi cichlid fishes, which was published in Malinsky *et al.* (2018); 2) a small simulation dataset comprising 20 species and 20Mb of sequence generated using the `ms’` (Kelleher *et al.* 2016) software; 3) a large simulation dataset with 100Mb of sequence and 100 species. In the simulated data, directional admixture events were simulated at randomly selected time points, with uniform distribution between the initial split time and the present, between a randomly selected pair of branches existing at that time point, and with admixture proportions drawn from a beta distribution rescaled to be between 0% and 30% with a maximum around 5% to 10%. Diploid samples were produced by combining two independently simulated haploid sequences. The outgroup was defined as diverging from other species two million generations before present and having effective population size of one (N_e_ = 1) to ensure all differences are fixed in the outgroup. The common ancestor of all the other species was then set to be at one million generations ago. Further details about the datasets and the parameters used in the simulations are outlined in Table 1.

**Table 1.** An outline of datasets used to evaluate the performance of Dsuite.

| Dataset | Species | Samples | Trios | Sequence length | SNPs | Simulation parameters |  |  |  |
| --- | --- | --- | --- | --- | --- | --- | --- | --- | --- |
| | | | | | | $\mu, \rho^*$<br>($10^{-8}$) | $N_e$<br>($10^3$) | Gene flow events | Age (generations) |
| Malawi scaffold_0 | 73 | 131 | 62,196 | 16Mb | 612,889 | ----- | Real data | ----- |  |
| Simulation small | 20 | 40 | 1,140 | 20Mb | 4,342,771 | 1 | 50 | 5 | 1 million |
| Simulation large | 100 | 200 | 161,700 | 100Mb | 97,201,601 | 1 | 50 | 10 | 1 million |

\* $\mu$ - per generation mutation rate; $\rho$ - per-generation recombination rate

### Computational efficiency

To assess computational efficiency of Dsuite, we calculated the D statistics for all combinations of trios with three other software packages: ADMIXTOOLS, HyDe, and Comp-D. For the Malawi cichlids and for the large simulated datasets, Dsuite was the most efficient of the programs in terms of both memory requirements and run time. For the small simulated dataset, Dsuite was still the most memory efficient, but ADMIXTOOLS and HyDe were faster. The full results are shown in Table 2.

**Table 2.** A comparison of Dsuite and a number of other tools in terms of computational efficiency of D statistic estimation.

| Dataset | Software | Options | Peak memory | Run time |
| --- | --- | --- | --- | --- |
| Malawi scaffold_0 | Dsuite Dtrios | --no-f4-ratio | 92MB | 74m59s |
|  | Admixtools qpDstat | blgsize: 0.01 | 27,212MB | 125m2s |
|  | HyDe run_hyde.py | none | 178MB | 231m38s |
|  | Comp-D* | -d -H -b10 | 8,300MB | 24hours+ |
| Simulation small<br>(20 species) | Dsuite Dtrios | --no-f4-ratio | 8MB | 28m18s |
|  | Admixtools qpDstat | blgsize: 0.01 | 17,100MB | 13m59s |
|  | HyDe run_hyde.py | none | 258MB | 19m38s |
|  | Comp-D* | -d -H -b10 | 22,100MB | 24hours+ |
| Simulation large<br>(100 species) | Dsuite Dtrios | --no-f4-ratio | 223MB | 215m52s (×100§) |
|  | Admixtools qpDstat | blgsize: 0.05 | 1,117,314MB | 331m39s (×100§) |
|  | HyDe run_hyde.py | none | 18,716MB | 576m32s (×100§) |
|  | Comp-D* | -d -H -b10 | 1,000,185MB+ | 24hours+ (×100§) |

\*Comp-D cannot use allele frequencies calculated across multiple individuals, so only one individual per species included.
§Because of the size of the dataset, we divided the analysis into 100 equally sized jobs to run in parallel; the run time and memory requirements are given for the first job

The advantage of the small memory footprint of Dsuite was most pronounced in the analysis of the large simulation dataset. There, ADMIXTOOLS and Comp-D required over 1 Terabyte of RAM and HyDe over 18 Gigabytes, while the Dsuite run required less than 223MB. The difference in memory efficiency between Dsuite and especially ADMIXTOOLS and Comp-D remained more than two orders of magnitude also for the two other datasets. In terms of speed, Comp-D stood out as being substantially slower. We cancelled all the Comp-D runs after 24hours with only a small proportion of the trios completed. Among Dsuite, ADMIXTOOLS, and HyDe, the run time differences were up to ∼2-3 fold depending on the dataset (Table 2).

While the Dsuite analysis was run directly on the VCF file, all other software required format conversion. For ADMIXTOOLS, we first obtained data in the PED format using VCFtools v0.1.12b (Danecek *et al.* 2011) with the `--plink` option, and then translated these into the software-specific EIGENSTRAT format using the `convertf` program, which is included in the ADMIXTOOLS package. Data conversion into the PHYLIP input format for HyDe and Comp-D was done using the vcf2phylip script (Ortiz 2019). The additional run and set-up time needed for these conversions was excluded from the run times shown in Table 2.

## Results and interpretation

In this section we use the small simulated dataset to illustrate the outputs of Dsuite and some topics related to the interpretation of the results. The results for the Malawi cichlid dataset are discussed in Malinsky *et al.* (2018).

We found tens of differences among the trio arrangements in the three output files produced by Dsuite `Dtrios` (Fig. 1A). The “BBAA” trio arrangements differed from the correct tree in 39 cases (3.4% of the trios), which illustrates that sister species do not always share the most derived alleles in the presence of gene flow, even in the absence of rate variation. However, unlike for the simulation, the correct tree is not known for most real-world datasets and the frequency of the “BBAA” pattern may then be a useful guide regarding the population relationships. The “Dmin” arrangements differed from the correct tree in 124 trios (10.9%).

**Figure 1:**
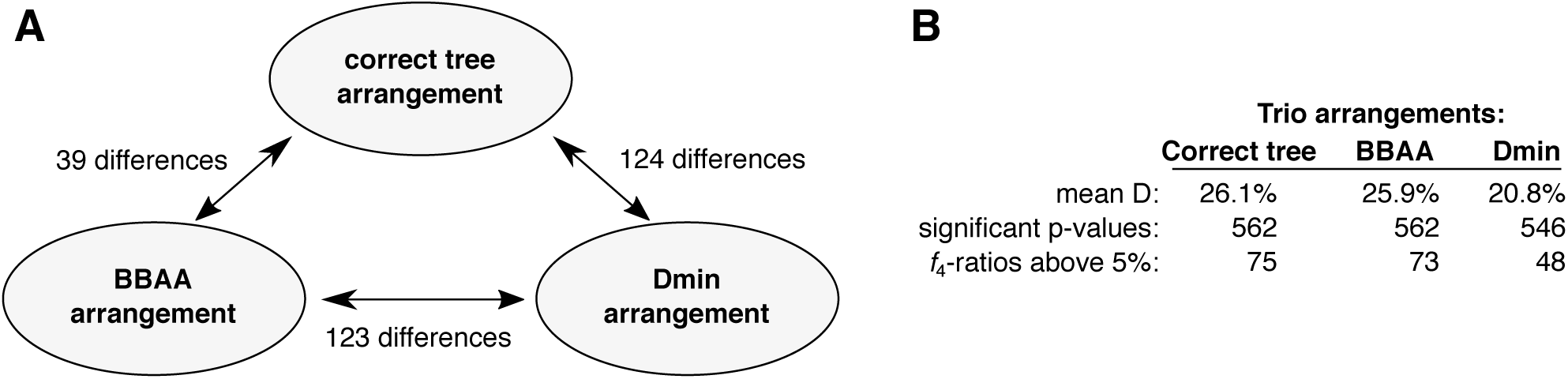
Summary of Dtrios output for the small simulated dataset (20 species, 1,140 trios, 5 gene-flow events). **A)** The number of differences in trio arrangements between the three different output files. **B)** A brief summary comparing the results with the three alternative arrangements.

Keeping in mind that only five gene flow events were simulated, it is notable that almost half of the D statistics were significantly elevated, e.g. 546 (47.9%) even in the “Dmin” arrangement which provides a lower bound on the D value for each trio (Fig. 1B). Using the *f*_4_-ratio measure, we found that admixture proportions above 5% were estimated for at least 48 trios. This demonstrates that D and *f*_4_-ratio statistics are correlated and that a significantly elevated result for a trio does not necessarily pinpoint the pair of populations involved in a gene flow event.

The tree in Fig. 2 shows the true simulated relationships between the 20 species together with the five gene flow events and their admixture proportions. The output of Dsuite `Fbranch` inference is then plotted in the inset heatmap, revealing how the f-branch statistic is useful in guiding the interpretation of correlated *f*_4_-ratio results. Ten out of the 568 f-branch (*f*_*b*_) signals are stronger than 5%, much fewer than the 73 signals identified from the raw trio analysis with the “BBAA” trio arrangements.

**Figure 2:**
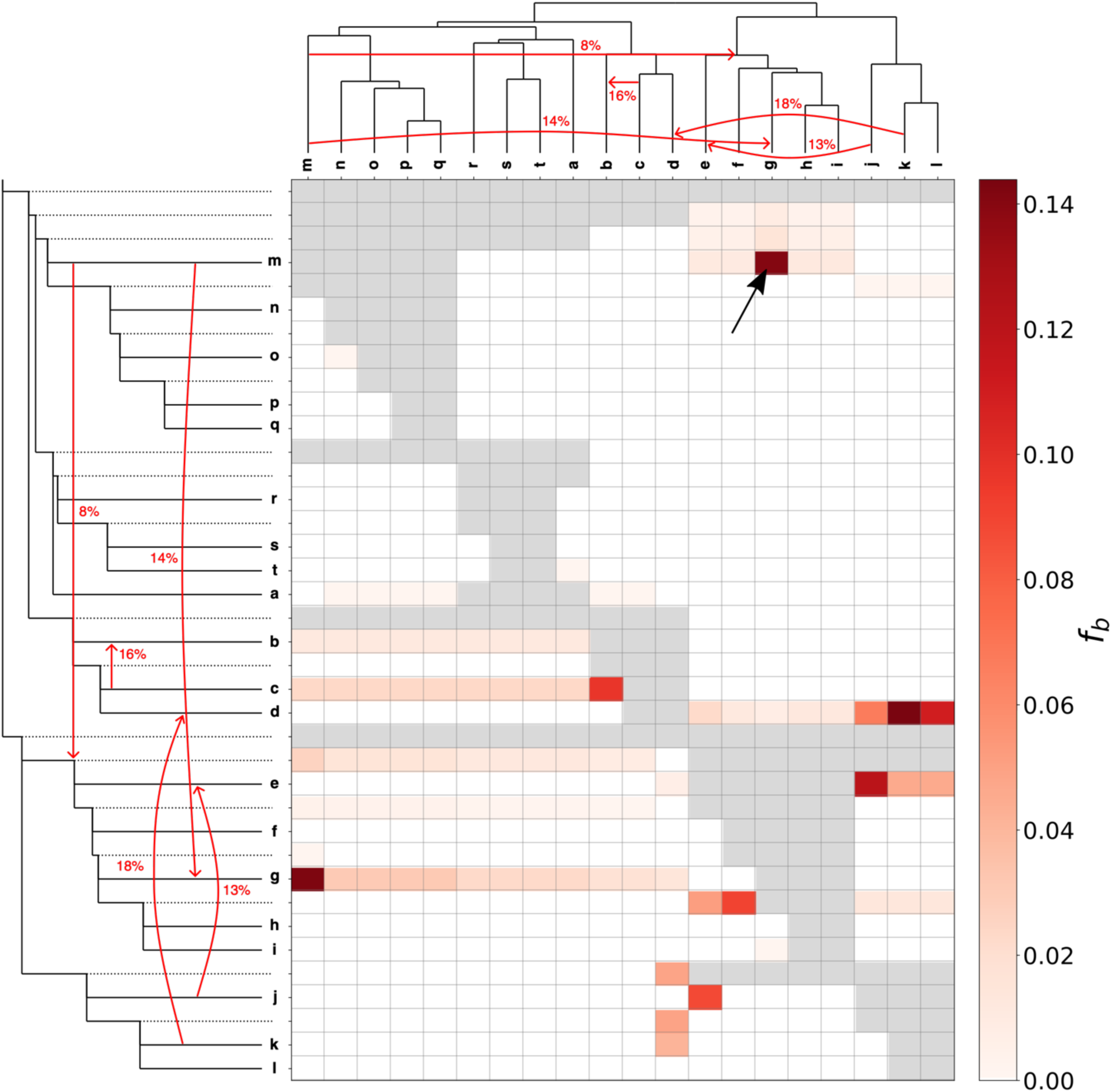
Fbranch results for the small simulated dataset. The tree used for simulating the data is shown along the x and y axes (in ‘laddered’ form along the y axis), together with the simulated gene-flow events and true admixture proportions. The matrix shows the inferred f-branch statistics, showing excess allele sharing between the branch of the ‘laddered’ tree on the y axis (relative to its sister branch) and the species identified on the x axis. As an example, the cell highlighted by the black arrow refers to excess allele sharing between species g and the branch leading to species m, relative to its sister, the internal branch above species n, o, p, and q.

The reduction of information and the visualization provided by f-branch facilitates narrowing down the number of possible acceptor and donor lineages involved in a gene flow event and should be seen as an aid for formulating specific gene flow hypotheses in a large data set that can be followed up individually by other methods, for example in a model-based inference framework by software such as fastsimcoal2 (Excoffier *et al.* 2013). In particular, the ten f-branch signals stronger than 5% correctly identify seven out of the nine branches involved in gene flow events. Six of these signals correctly pinpoint both branches involved in gene flow events ((d, k), (e, j), (m, g), (c, b)). However, a single gene flow event between two branches can still produce more than one f-branch signal. For example, the gene flow event from m into g above produces elevated values for both f_b=g_(C=m), i.e. the branch leading to g and species m, and its ‘mirror image’ f_b=m_(C=g), branch leading to m and species g. Furthermore, the gene flow from m into g produces correlated signals between g and lineages related to m (e.g. n, o, p, q) because of the shared ancestry between these lineages and m. This generally manifests in horizontal lines of correlated signals in the f-branch plots as shown in Fig. 2. Finally, note that an f-branch result in itself does not indicate directionality of gene flow. We suggest using 5-taxon tests, when possible, for inferring directionality (Pease & Hahn 2015; Svardal *et al.* 2019).

## Discussion

The Dsuite software package brings together a number of statistics for learning about admixture history from patterns of allele sharing across populations or closely related species. In particular, by being computationally efficient, it facilitates the calculation of the *D* and *f*_4_-ratio statistics across tens or even hundreds of populations, meeting the needs of ever growing genomic datasets. Correct interpretation of the results of a system of *D* and *f*_4_-ratio tests remains challenging and is an active area of research. In real datasets, imbalances in allele sharing that lead to significantly elevated D and *f*_4_-ratio statistics can result from specific scenarios involving ancestral population structure (Durand *et al.* 2011; Eriksson & Manica 2012) and variation in substitution rates (Pease & Hahn 2015). Even when all allele sharing imbalances are caused by introgression more work remains to be done to reliably pinpoint all introgression events and infer the networks of gene flow that may characterise relationships between many populations or closely related species. Dsuite implements tools that aid the interpretation of the results, including the f_d_ and f_dM_ statistics suited for applying to genomic windows and the f-branch statistic which aids in assigning the gene flow to particular branches on the population or species tree.

## Acknowledgements

We would like to thank Richard Durbin and Walter Salzburger for useful discussions and comments.

## Author contributions

MilMal developed the Dsuite software package with assistance from MicMat regarding tree-based operations, HS conceived the f-branch statistics and coded the plotting function for it. All authors contributed to and approved the manuscript.

## Data availability

The Malawi cichlid data and the simulated data used in this manuscript are available through the Dsuite GitHub repository (https://github.com/millanek/Dsuite).

